# An improved method for culturing myotubes on laminins for the robust clustering of postsynaptic machinery

**DOI:** 10.1101/664268

**Authors:** Marcin Pęziński, Patrycja Daszczuk, Bhola Shankar Pradhan, Hanns Lochmüller, Tomasz J. Prószyński

## Abstract

Motor neurons form specialized synapses with skeletal muscle fibers, called neuromuscular junctions (NMJs). Cultured myotubes are used as a simplified *in vitro* system to study the postsynaptic specialization of muscles. The stimulation of myotubes with the glycoprotein agrin or laminin-111 induces the clustering of postsynaptic machinery that contains acetylcholine receptors (AChRs). When myotubes are grown on laminin-coated surfaces, AChR clusters undergo developmental remodeling to form topologically complex structures that resemble mature NMJs. Needing further exploration are the molecular processes that govern AChR cluster assembly and its developmental maturation. Here, we describe an improved protocol for culturing muscle cells to promote the formation of complex AChR clusters. We screened various laminin isoforms and showed that laminin-221 was the most potent for inducing AChR clusters, whereas laminin-121, laminin-211, and laminin-221 afforded the highest percentages of topologically complex assemblies. Human primary myotubes that were formed by myoblasts obtained from patient biopsies also assembled AChR clusters that underwent remodeling *in vitro*. Collectively, these results demonstrate an advancement of culturing myotubes that can facilitate high-throughput screening for potential therapeutic targets for neuromuscular disorders.

## Introduction

Vertebrate neuromuscular junctions (NMJs) are synapses between motor neurons and skeletal muscle fibers. The function of this type of synapse is to transmit signals from the central nervous system to muscles and thus stimulate their contraction. The nerve terminal releases the neurotransmitter acetylcholine (ACh), which binds to postsynaptic ACh receptors (AChRs) that are located on the surface of muscle fibers. For efficient synaptic transmission, muscle fibers need to accumulate a high density of AChRs in their postsynaptic membrane ^1, 2, 3, 4^. Muscles cluster components of postsynaptic machinery around day 12 of embryonic development (i.e., before motor neuron axons approach muscle fibers)^5, 6^. Innervation leads to the dispersion of preexisting AChR clusters^2, 7^. A single postsynaptic machinery is formed per muscle fiber directly below the nerve terminal. The nerve plays a crucial role in organizing NMJ postsynaptic machinery by secreting signaling molecules, such as agrin and ACh, which regulate the clustering and dispersion of AChRs, respectively (9, 10, 12 Lin mei review, Sanes Agrin counterbalance ACH secretion ^2, 8-11^. Nerve-derived agrin binds to the surface of low-density lipoprotein receptor-related protein (Lrp4), which activates muscle-specific kinase (MuSK)^12^. This triggers an intracellular signaling cascade that activates AChR clustering by the scaffold protein rapsyn^2^. In the first postnatal weeks, NMJs grow in size and undergo developmental remodeling from simple plaque-shaped structures to topologically complex assemblies^2, 13^. During this process, postsynaptic machinery becomes perforated with scattered openings that, with time, become more numerous and fuse with each other to form indentations between AChR-rich branches that transform synapses into pretzel-like shapes^14,15^. The molecular mechanisms that underlie developmental remodeling are poorly understood.

The clustering process and synapse remodeling are facilitated by macromolecular complexes that are involved in cellular adhesion that interact with extracellular matrix (ECM) components and the cytoskeleton. The major ECM receptors contain integrin complexes and the dystrophin-associated glycoprotein complex^16-18^. These protein assemblies stabilize postsynaptic components and provide a platform for the recruitment of signaling molecules that regulate postsynaptic specialization^17, 19^. Muscle cells form a thick ECM around the fiber that contains various laminins, collagens, fibronectin, and other glycoproteins^20^. Laminins are heterotrimeric glycoproteins that are composed of α, β, and γ chains that are encoded by different genes^21, 22^. The different laminin trimers have various specificities for cellular receptors and can regulate different signaling pathways^23-26^. The basal lamina (BL) at the synaptic cleft has a specific molecular composition that contains laminin α4, α5, and β2 isoforms that are mostly absent in extrasynaptic regions of muscle fibers^21, 27^. These ECM components are crucial for the proper development of NMJs. Laminin β2 knock-out mice have severe phenotypes of reduced synaptic folding and drastically decreased synaptic transmission^28^. In laminin α4/α5 double-mutant mice, NMJs form normally but fail to undergo developmental remodeling^29^. A similar phenotype was observed with the muscle-specific deletion of dystroglycan, the laminin receptor in muscle cells^29^. This indicates that the laminin-dystroglycan interaction is crucial for regulating NMJ developmental remodeling. Studies of postnatal NMJ reorganization *in vivo* are complicated by the fact that different cells (e.g., motor neurons and Schwann cells) may contribute to this process. Aneurally cultured differentiated myotubes are often used as a simplified model to study postsynaptic machinery^30-32^. The stimulation of cultured myotubes with agrin is the most commonly used method to induce the formation of AChR clusters *in vitro*^31, 32^. The major limitation of this approach, however, is that AChR clusters do not undergo the remodeling process. The formation of AChR clusters in cultured myotubes can also be stimulated by laminins^30, 33^. Recent studies demonstrated that the culturing of muscle cells on laminin-111-coated surfaces leads to the formation of topologically complex clusters of postsynaptic machinery that resemble clusters at the NMJ *in vivo*^*30*^. Moreover, the complex shape of the cluster is achieved through morphological transformation that is similar to such transformation at the NMJ and involves the formation of perforations in the initially plaque-shaped clusters. Cultured myotubes that were derived from primary myoblasts or C2C12 myoblasts were shown to utilize podosomes to remodel postsynaptic machinery^34, 35^. These actin-rich organelles are involved in cellular adhesion and the degradation of ECM components, including laminins ^36, 37^. Little is known about the mechanisms of postsynaptic machinery remodeling *in vivo*. Myotubes cultured on laminin are the only *in vitro* system where the AChR clusters undergo developmental remodeling, providing the model to study the underlying mechanisms.

Myotubes that form AChR clusters can be used to study the mechanisms that underlie pathological processes in both congenital and autoimmune muscle disorders ^38, 39^. Defects in NMJ remodeling from “plaque” to “pretzel” shapes are frequently observed in many models of neuromuscular disorders^15, 40^. Some of these disorders still have an unknown etiology, thus highlighting the importance of studying the mechanisms of NMJ development.

The present study demonstrated an improved protocol for culturing myoblasts that allowed the reproducible formation of AChR clusters. We tested several commercially available laminin isoforms for their ability to induce AChR cluster formation and their ability to promote AChR cluster remodeling. Our observations provide an important basis for high-throughput genetic screening and potential drug development. We also found that human primary myotubes that were derived from myoblasts obtained from patient biopsies were also able to form AChR clusters with complex topology that contained synaptic podosomes.

## Results

### Optimal culturing of myoblasts for efficient AChR clustering

The postsynaptic machinery at NMJs is a topologically complex structure that contacts hundreds of different proteins (Fig. 1A). Studying postsynaptic machinery organization *in vivo* in tissues can be experimentally challenging. Differentiated myotubes that are cultured *in vitro* provide a simplified model to study basic processes that underlie postsynaptic specialization^30,32^. Three methods are the most commonly used to stimulate myotubes for AChR cluster formation. The glycoprotein agrin can be added to the media or deposited at the culturing surface^31, 32^. Soluble agrin triggered the formation of small AChR aggregates at apical and lateral surfaces of myotubes (Fig. 1B), whereas precoated agrin induced the formation of much larger oval assemblies (Fig. 1C)^32^. Alternatively, myotubes can be stimulated for AChR clustering by precoating the culture surface with laminins^30^. Myotubes derived from C2C12 cells or primary myoblasts that were cultured on laminin formed numerous large AChR clusters at the bottom of myotubes that often acquired a complex topology that was reminiscent of postsynaptic machinery at the NMJ (Fig. 1D)^30, 41, 42^

**Figure 1.**
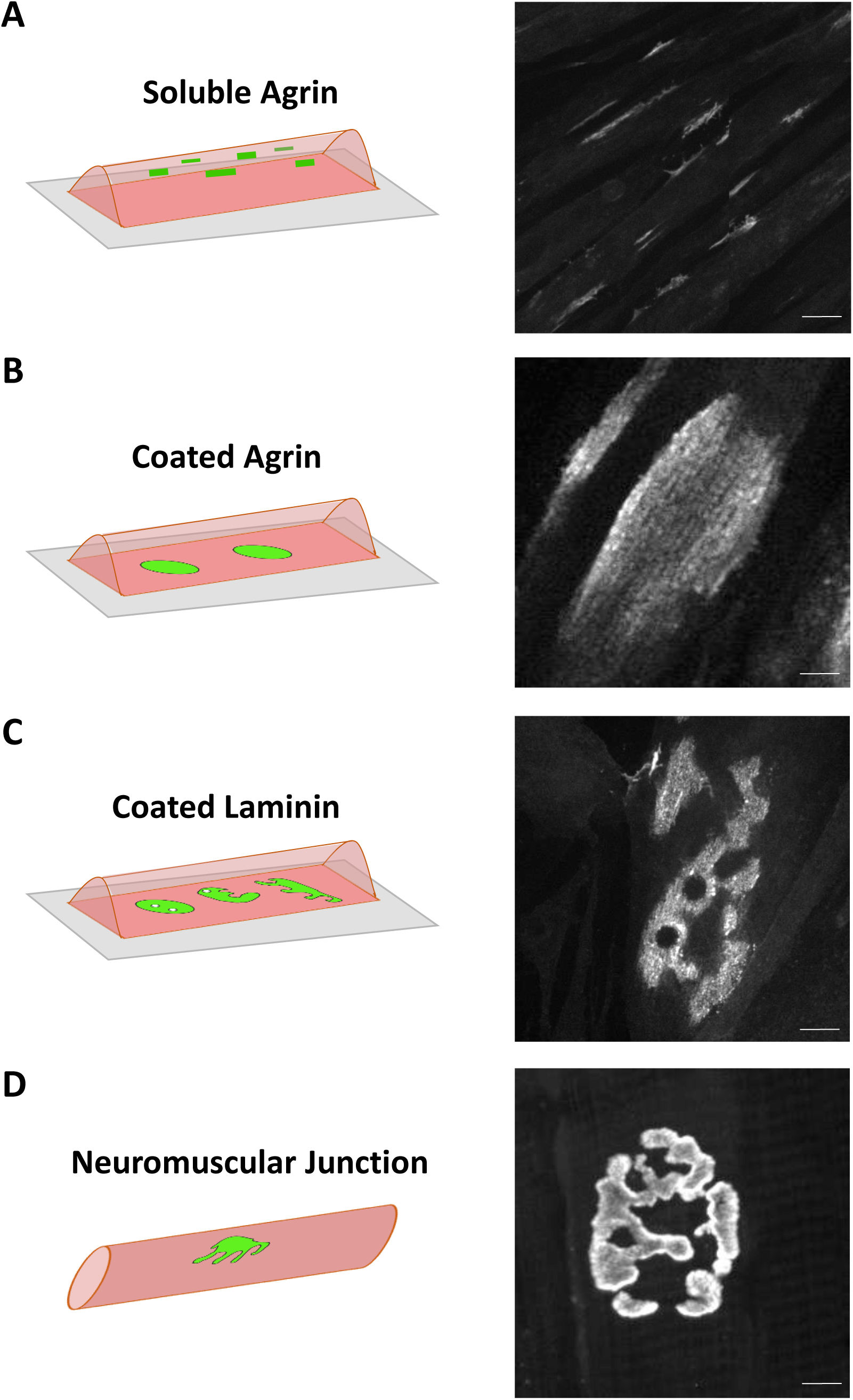
Models for studying muscle postsynaptic machinery. (**A**) Differentiated C2C12 myotubes that were stimulated with a neuron-specific splice variant of the glycoprotein agrin that was supplemented to the media formed small, elongated AChR clusters on the lateral and apical sides of the myotubes. (**B**) C2C12 myotubes that were grown on a laminin substratum with deposited agrin patches formed large uniform AChR assemblies at the basal surface of the myotubes. (**C**) C2C12 myotubes that were grown on laminin-111-coated surfaces formed clusters of postsynaptic machinery that underwent developmental remodeling to topologically complex assemblies that resembled the shape of postsynaptic machinery at the NMJ. (**D**) Complex structure of postsynaptic machinery at the murine NMJ. Scale bar = 25 μm in A and 6 μm in B-D.

Laminin-cultured myotubes are particularly useful because they can provide insights into the mechanism of postsynaptic machinery remodeling. This system has been used by many laboratories because it is relatively easy. Nonetheless, some groups have experienced problems with obtaining a reproducibly high number of AChR clusters that is sufficient for downstream experiments. Our laboratory found that most problems with laminin-cultured myotubes result from the inappropriate culturing of cells before dedifferentiation. The most common source of such problems is the culturing of myoblasts at an inappropriate density. The present study reports a relatively simple method of myoblast culturing that yields the most reliable results. Fig. 2A shows images of the optimal density for cell splitting (green frame) and cells that were grown too densely or too sparsely (red frames). Myoblasts that are too sparse in cultures grow slowly and tend to form clusters. Conversely, excessive cell-cell contacts lead to dedifferentiation and the loss of myogenic potential. In such case, the cells can still form elaborate myotubes but fail to cluster AChRs or the clusters are too small, underdeveloped, or scarce. Consequently, every subsequent generation of myoblasts has a lower capacity to form numerous and mature AChR clusters. For optimal experimental reproducibility, myoblasts should be of a low passage number and should be analyzed after the same number of amplifications. Fig. 2B shows an optimal scheme of passaging that we implement in our laboratory to obtain a large number of cells (up to 1000 vials) at passage 5 (P5). We do not use cells that are amplified for more than five passages in our experiments.

**Figure 2.**
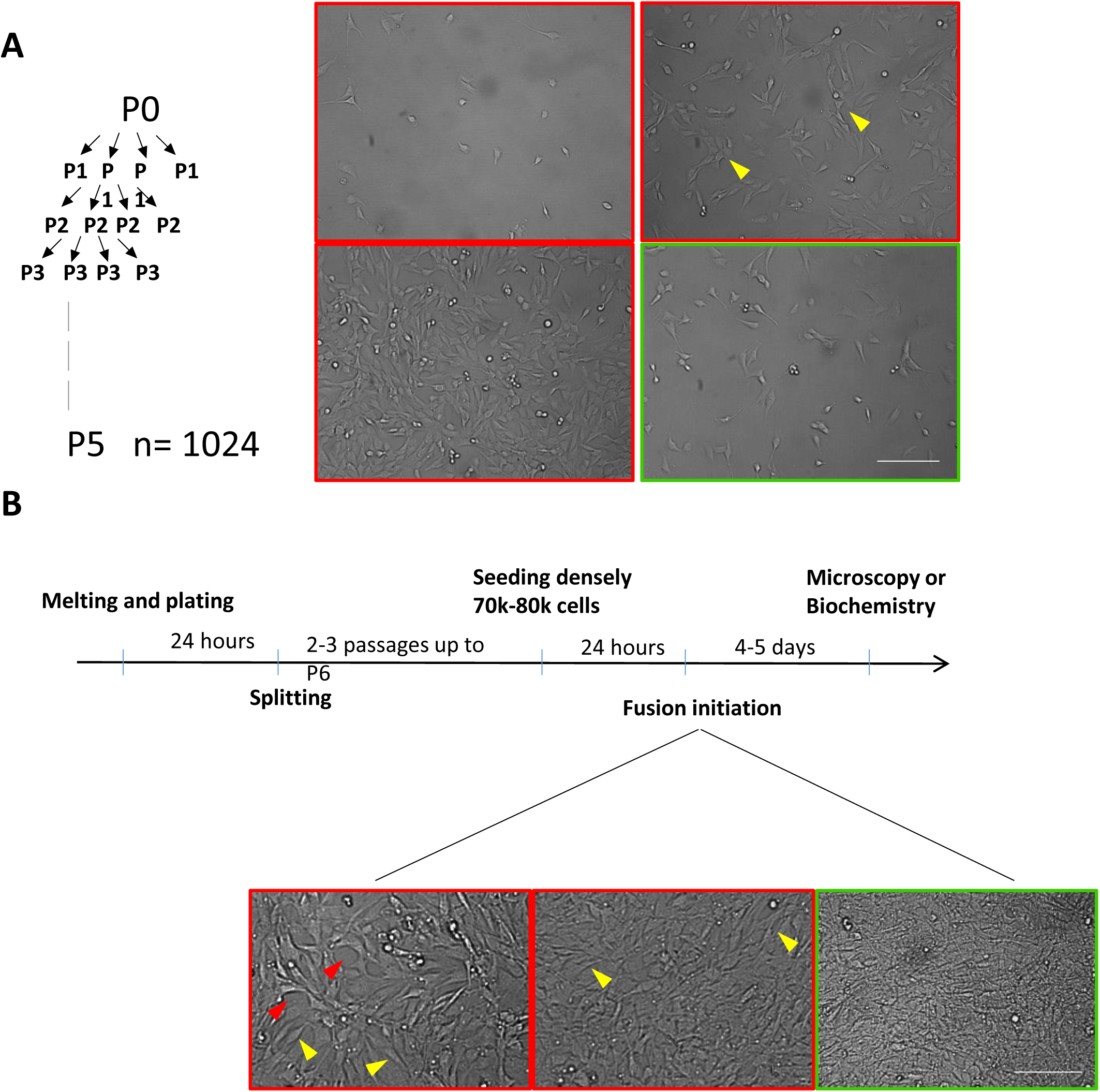
Method for culturing of C2C12 cells to obtain high yields of AChR clusters. (**A**) Schematic diagram of C2C12 cell passages to obtain a high number of cell stocks at an equally low number of passages. (**B**) Examples of suboptimal (red boxes) and optimal (green box) cell densities for passaging. Yellow arrows show the area of excessive cell contacts that should be avoided. (**C**) Schematic diagram of the key steps during myotube differentiation that are important for obtaining high yields of AChR clusters. The images on the bottom show optimal (green box) and suboptimal (red boxes) cell densities for fusion initiation. Red arrowheads show areas that are not covered by cells. Yellow arrows show regions where single-cell morphology can be observed. For the highest amount of fully differentiated myotubes with AChR clusters, myoblasts should form a dense cell layer in which cell morphology is difficult to distinguish. Scale bar = 600 μm in B and 450 μm in C.

The next critical step for obtaining a large number of AChR clusters in cultures involves performing appropriate steps for cell preparation for differentiation on laminin-coated surfaces. After thawing, frozen cells should be passaged twice (up to P7) and seeded on laminin in sufficient density (1.1 × 10^5^ cells/cm^2^) for fusion between 24 and 48 h. The prolongation of cell growth on laminin could result in substrate degradation and a poor yield of AChR clusters. Obtaining a high density of cells upon differentiation induction is critical (Fig. 2C) because a culture that is too sparse results in poorly differentiated myotubes and myoblasts that are too dense and lose their myogenic potential^43, 44^. Finally, differentiating myotubes should not be agitated during the fusion process. Larger myotubes tend to detach easily during medium washes, which should be avoided. Rapid shifts in temperature also result in myotube detachment. Therefore, we do not recommend removing differentiating cells from the incubator during the fusion process.

### AChR clustering in C2C12 myotubes that are cultured on different laminins

Laminin-111 has been routinely used to induce AChR clustering^30^. Various laminin isoforms have different properties. We examined the ability of different laminin isoforms to induce the clustering of muscle postsynaptic machinery in cultured C2C12 myotubes. We cultured cells on laminin-111, laminin-121, laminin-211, laminin-221, laminin-411, laminin-421, laminin-511, and laminin-521. Coating the culture surfaces with laminin-111, laminin-121, laminin-211, laminin-221, laminin-511, and laminin-521 led to the formation of numerous AChR clusters (Fig. 3A). Surprisingly, the commonly used laminin-111 reproducibly yielded a lower number of clusters compared with the other laminins (Fig. 3B). In contrast, laminin trimers that contained the α4 chain (i.e., laminin-411 and laminin-421) were very inefficient in inducing AChR aggregates (Fig. 3A, B), and AChR aggregates that formed had a fragmented appearance (Fig. 3A, insets). This could be because of the lack of N-terminal domain in the α4 chain, which is present in other α isoforms and which serves as a binding site for integrin and other cell surface receptors (Fig. 3D). To exclude the possibility that the observed differences were attributable to varying levels of laminins that adhered to the surfaces during coating, we analyzed material that adhered to the culture wells after coating using silver staining. As shown in Supplementary Fig. S1, similar amounts of each laminin were recovered in three independent experiments. The culturing of myotubes on various laminins led to the similar induction of AChR production, analyzed by Western blot (Fig. 3B). Thus, the differences that we observed likely reflected the ability of individual laminins to induce AChR clustering.

**Figure 3.**
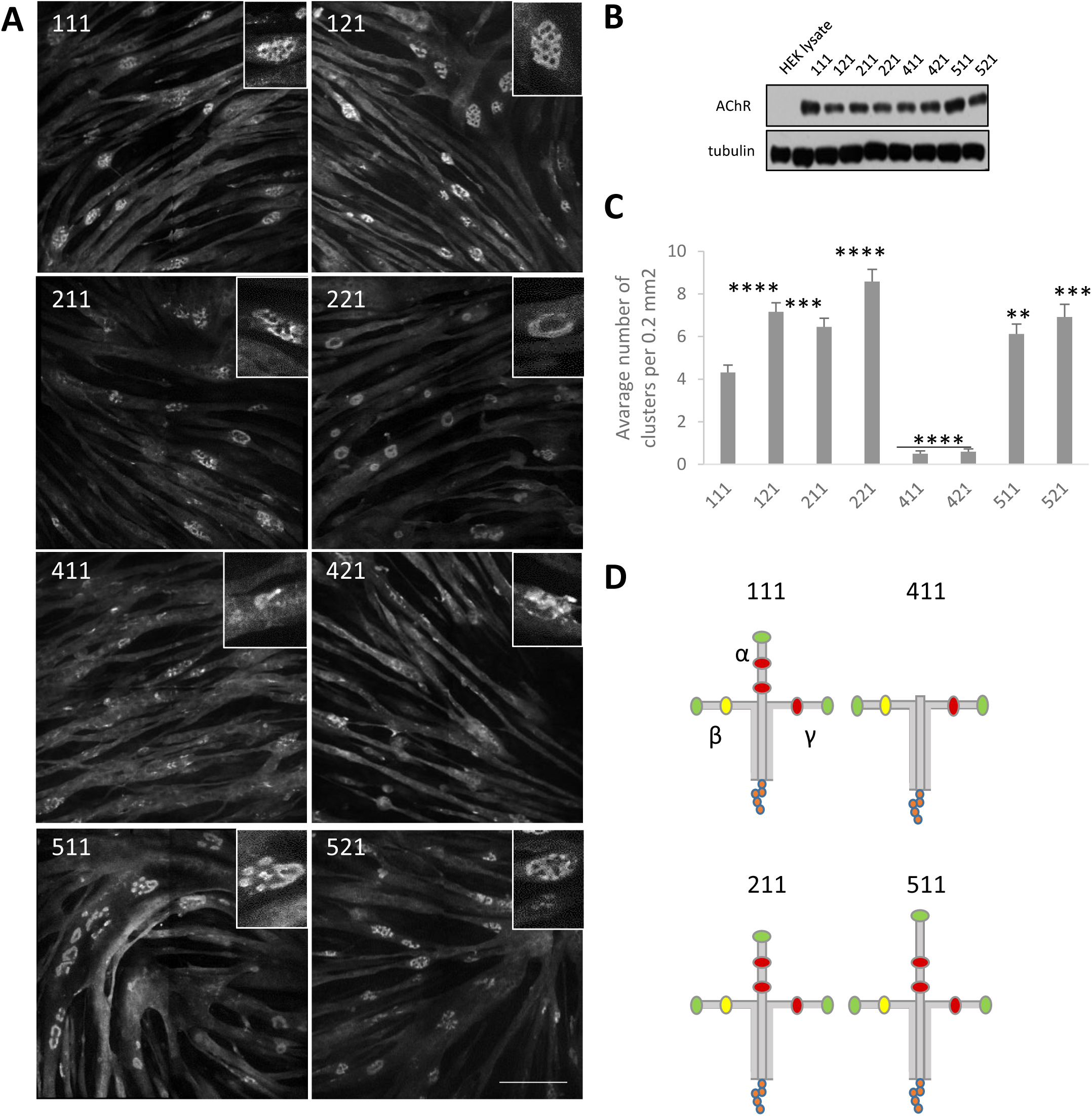
Clustering of AChRs in C2C12 myotubes cultured on different laminins. (**A**) Representative images of AChR clusters in myotubes that were cultured on the indicated laminins. AChRs were visualized with α-bungarotoxin. Scale bar = 150 μm. (**B**) Different laminins induced similar levels of AChR expression in myotubes. HEK293 cell lysates were used as a negative control. Tubulin was used as a loading control. (**C**) Quantification of AChR clusters formed by myotubes that were grown on the indicated laminins. Quantifications represent the average number of clusters. For all laminins except laminin-111, laminin-411, and laminin-421, the total number of clusters that were used for the analysis was > 300. For laminin-111, 216 clusters were used. For laminin-411, 25 clusters were used. For laminin-421, 30 clusters were used. The clusters were collected from five independent experiments. \**p* < 0.05, \*\**p* < 0.005, \*\*\**p* < 0.005, **** *p* < 0.0005, compared with laminin-111 (*t*-test). (**D**) Schematic representation of laminin-111, 211, 411, 511. LN domains are shown in green, L4 domains in red, LF domain in yellow and LG domains in orange.

### Effect of different laminins on AChR cluster formation in C2C12 myotubes

The assemblies of postsynaptic machinery in laminin-cultured myotubes underwent developmental remodeling that resembled maturation of the NMJ *in vivo* (Fig. 4A)^30, 45^. During this process, oval, plaque-shaped AChR clusters (Fig. 4A, C) became perforated and acquired a much more complex topology. Some of the remodeled clusters acquired C-shaped topology, in which a large portion of AChRs was removed from the center of the assembly (Fig. 4A, E)^30^. Other clusters had a perforated pretzel-like shape with actin-rich structures, known as synaptic podosomes, that interdigitated AChR-rich domains (Fig. 4A, G)^34, 45^.

**Figure 4.**
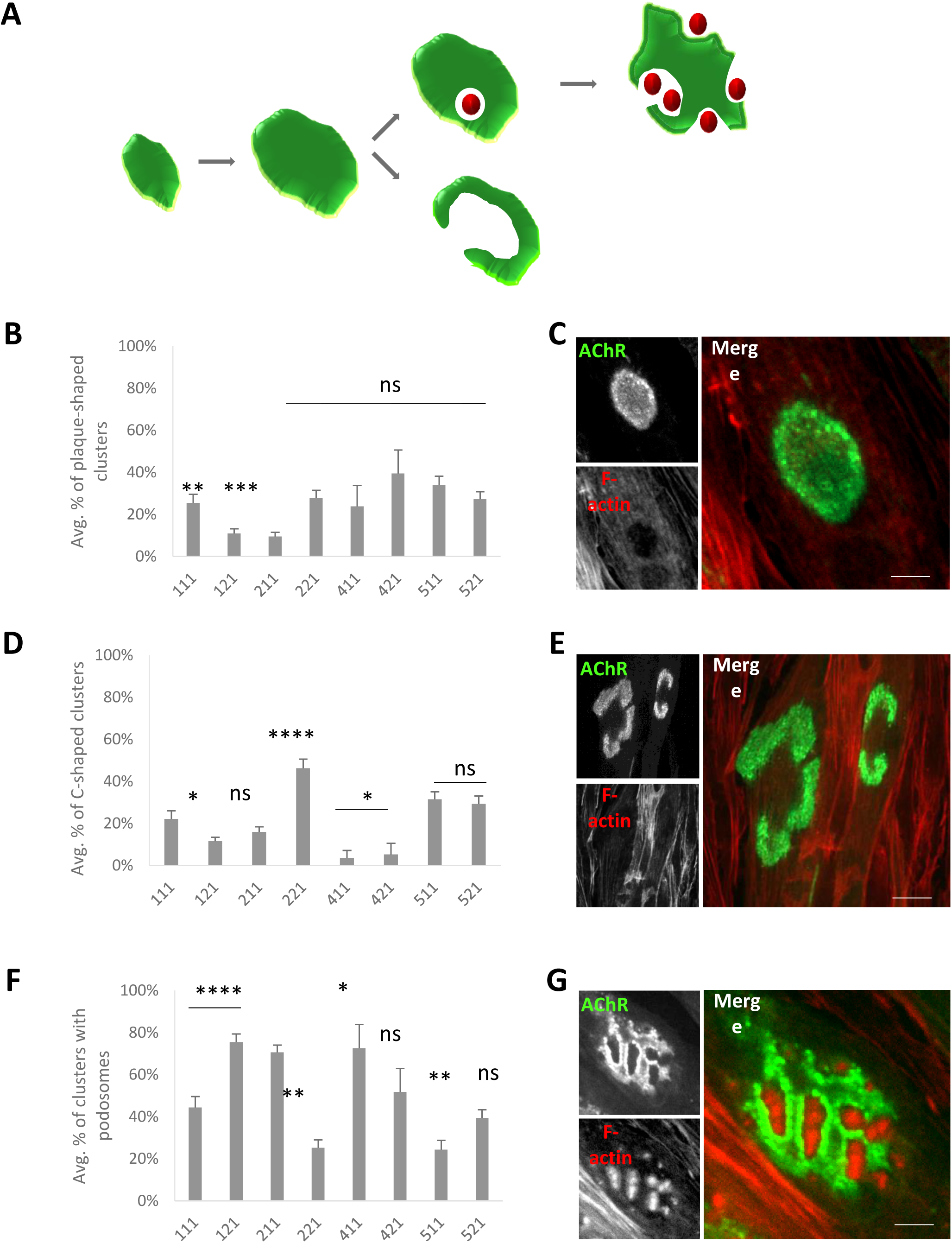
Different AChR cluster morphologies in C2C12 myotubes grown on various laminins. (**A**) Schematic diagram showing developmental remodeling of postsynaptic machinery. Actin-rich podosomes are represented as red spheres. (**B**) Quantification of plaque-shaped clusters, presented as the average percentage of clusters. (**C**) Example of plaque-shaped cluster. (**D**) Quantification of C-shaped clusters, presented as the average percentage of clusters. (**E**) Example of C-shaped cluster. (**F**) Quantification of perforated clusters that contained synaptic podosomes, presented as the average percentage of clusters. (**E**) Example of cluster that contained actin-rich podosomes. AChR (green) was visualized with α- bungarotoxin. F-actin (red) was visualized with phalloidin. Scale bar = 6 μm. Quantifications represent the average number of clusters. For all laminins except laminin-111, laminin-411, and laminin-421, the total number of clusters that were used for the analysis was > 300. For laminin-111, 216 clusters were used. For laminin-411, 25 clusters were used. For laminin-421, 30 clusters were used. Clusters were collected from five independent experiments. \**p* < 0.05,\*\**p* < 0.005, \*\*\**p* < 0.005, **** *p* < 0.0005, compared with laminin-111 (*t*-test).

Interestingly, C2C12 myotubes that were cultured on different laminins exhibited alterations of the maturation of AChR clusters. Of the tested laminin isoforms, none increased plaque-shaped clusters compared with isoform-111 (Fig. 4B, C). Cultures on laminin-121 and laminin-211, which led to the formation of numerous clusters (Fig. 3A, B), produced significantly fewer immature plaque-like AChR aggregates (Fig. 4B, C). Laminin-121 and laminin-211 appeared to induce fewer C-shaped clusters than laminin-111, although the effect of laminin-211 did not reach statistical significance (Fig. 4D). The highest percentage of C-shaped structures was observed in myotubes that were cultured on laminin-221 (Fig. 4D, E). Laminin-221, similar to laminin-511, induced significantly fewer perforated clusters with actin-rich podosomes compared with laminin-111 (Fig. 4F, G). The highest percentage of perforated clusters (> 70%) was formed by cells that were grown on laminin-121 and laminin-211 (Fig. 4F, G). Interpreting the types of clusters formed by cells that were cultured on laminins that contained the α4 isoform (i.e., laminin-411 and laminin-421) was more difficult because these cells made hardly any aggregates (Fig. 3A, B). These findings indicate that different types of laminins that are used for coating exert differential effects on the developmental remodeling of AChR clusters in cultured C2C12 myotubes.

### AChR cluster formation and remodeling in primary human myotubes

Myotubes that are derived from human patients myoblasts can be used to study the molecular processes that underlie pathological alterations associated with neuromuscular disease and screening novel therapeutics^46^. We analyzed the ability of human primary myotubes to form clusters of postsynaptic machinery on different laminins. In contrast to C2C12 myotubes, most of the laminins that were used for coating did not significantly influence the efficiency of AChR clustering. One notable exception was laminin-221, which led to the formation of fewer clusters (Fig. 5A) compared to all other isoforms. Most of the AChR clusters that formed in human myotubes had a simple, plaque-like shape (Fig. 5B, C). However, some AChR clusters underwent topological reorganization, with visible perforations within the clusters (Fig. 5D, E). These perforations appeared to be similar to those in C2C12-derived myotubes (Fig. 1, Fig. 4G). Perforations in C2C12 clusters are formed by synaptic podosomes^34^. Therefore, we examined whether human myotubes also form these organelles during cluster remodeling. Immunocytochemical analysis revealed that perforated areas in AChR assemblies in human myotubes contained actin-rich cores^34, 36^. In typical podosomes, these actin-rich structures are surrounded by the cortex domain that is enriched in proteins that are associated with cellular adhesion^34, 36, 41, 47^. As expected, AChR clusters in human myotubes also had strong vinculin and LL5β immunoreactivity in the cortex-like domain that abutted actin-rich cores (Fig. 5D, E). Interestingly, most of the tested laminins produced more clusters with complex topology than laminin-111 (Fig. 5F). Thus, AChR clusters in human primary myotubes were able to undergo developmental remodeling, and this process involved podosome formation.

**Figure 5.**
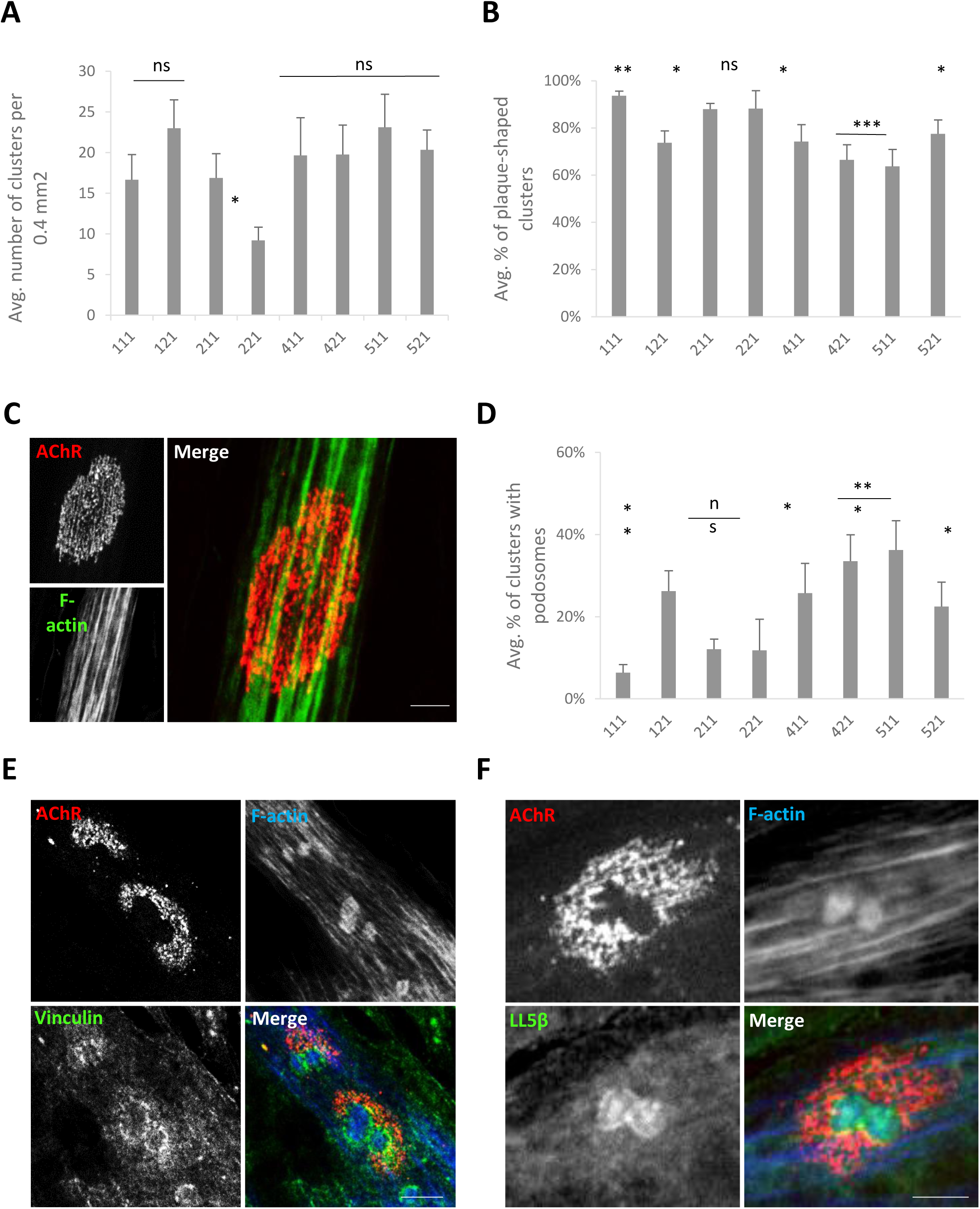
Effect of various laminins on AChR cluster formation and remodeling in human primary myotubes. (**A**) Quantification of AChR clusters formed by human primary myotubes that were cultured on the indicated laminins. (**B**) Quantification of plaque-shaped AChR clusters formed by human primary myotubes that were cultured on the indicated laminins, presented as the average percentage of clusters. (**C**) Example of plaque-shaped cluster. (**D**) Quantification of perforated clusters that contained synaptic podosomes, presented as the average percentage of clusters. (**E, F**) Examples of clusters that contained actin-rich podosomes that were immunostained for podosome cortex protein markers vinculin (**E**) and ll5 β (**F**). AChR (red) was visualized with α-bungarotoxin. F-actin (blue) was visualized with phalloidin. Scale bar = 6 μm. Quantifications represent the average number of clusters. For all laminins except laminin-221, the total number of clusters that were used for the analysis was > 140. For laminin-221, 83 clusters were used. Clusters were collected from three independent experiments. \**p* < 0.05, \*\**p* < 0.005, \*\*\**p* < 0.005, **** *p* < 0.0005, compared with laminin-111 (*t*-test).

## Discussion

Myotubes that are cultured on laminin-coated surfaces provide a unique *in vitro* system whereby clusters of muscle postsynaptic machinery undergo robust developmental remodeling^29, 30, 34^. Several laboratories, however, have experienced difficulties in culturing myotubes that reproducibly form AChR clusters with complex topology. Herein, we describe an improved protocol for culturing C2C12 muscle cells and human primary myoblasts to study the mechanisms that govern postsynaptic machinery development and remodeling. Our protocol is useful for obtaining and freezing a large number of cell stocks and utilizing cells for experimentation with a constant passage number. This significantly increases experimental reproducibility. We tested several laminin isoforms that are commercially available and found that laminin-121, laminin-211, laminin-221, laminin-511, and laminin-521 induced significantly more AChR clusters in C2C12 myotubes than the commonly used laminin-111. Moreover, we found that clusters of postsynaptic machinery that were formed in C2C12 myotubes cultured on laminin-121 and laminin-221 were the most developed. Myotubes that were derived from human primary myoblasts obtained from human biopsies also formed AChR clusters that underwent developmental remodeling, and these complex clusters contained synaptic podosomes. In contrast to C2C12, laminin-421 and laminin-511 were the isoforms that promoted formation of the most podosome-containing AChR clusters in human primary myotubes.

The molecular mechanisms that underlie the developmental remodeling of NMJs and muscle postsynaptic machinery are poorly understood, likely because of the various cell types (e.g., muscle cells, Schwann cells, and motor neurons) that could contribute to this process. *In vitro*-cultured myotubes provide a minimalistic system whereby at least some aspects of AChR regulation can be studied^13^. C2C12 and mouse primary myotubes utilize podosomes to remodel postsynaptic machinery when cultured on laminin^34^. There is no evidence, however, of the presence of such organelles at NMJs *in vivo*. Several podosome-associated proteins have been shown to be present at the muscle postsynaptic specialization, and they may be involved in the remodeling process^41, 45^.

NMJ postnatal transformation from “plaques” to “pretzels” has been shown to depend on the laminin-dystroglycan signaling pathway^29^. Integrins are also involved in the organization and stabilization of postsynaptic machinery, but their direct roles in developmental synapse remodeling is unclear^48^. Laminin trimers contain several binding sites for their surface receptors. The N’-terminal portion of most of α isoforms (missing in laminin α4) interacts with integrins and globular domains located at the C-terminus of laminin α chains bind α-dystroglycan^23^ (Fig. 3D). Each α chain contains five laminin globular subunits (LG1-5)^26^. Depending on the isoform of the α chain, different modules are utilized for the interaction with α-dystroglycan^24^. Despite differences in the localization of interaction sites, they all share the common feature of being calcium-dependent. Interestingly, agrin also interacts with α-dystroglycan in a calcium-dependent manner through its own laminin globular module^25^. Our observation that laminins containing the α4 chain induce clustering of AChRs in human primary muscles but not in C2C12, hints toward possibility that the formation of AChR clusters in C2C12 cells depends more than in human cells on interactions with integrins through the N’-terminal domain. It is also a possibility that efficiency of laminin polymerization through their LN domains is different for each tested isoform. This could result in differences in the accessibility to surface receptors and altered stiffness of the culturing surface, which are factors that may influence the efficiency of AChR clustering and remodeling. Another interesting line of investigation would be to test the affinities of laminin globular domains to α-dystroglycan in our experimental system, which may provide insights into the mechanisms of AChR cluster assembly.

In addition to being an intriguing biological process, the postnatal reorganization of synapses in topologically complex structures appears to be affected in many models of neuromuscular disorders^2, 40, 49^. Aberrant synaptic maturation may be linked to various pathological processes^2, 15^. Several neuromuscular myopathies have an unknown etiology. This highlights the importance of studying the molecular aspects of NMJ development. The present method for culturing and stimulating myotubes to form and remodel AChR clusters may facilitate the identification of novel synaptic regulators. The high reproducibility of culturing and robust formation of AChR clusters are important prerequisites for establishing high-throughput screening. The present study found that human cells that were derived from patient muscle biopsies formed AChR clusters *in vitro* that underwent the remodeling process, thus demonstrating the potential utility of this methodology for further studies that seek to improve diagnoses of neuromuscular disorders and elucidate their underlying mechanisms.

## Materials and Methods

### Passaging C2C12 cells

C2C12 myoblasts (catalog no. 13K011, Sigma) were cultured on 10 cm gelatin-coated plates. The surface of the plate was coated with 0.2% gelatin in double-distilled H2O that was then discarded after a few minutes. The plate was allowed to dry for 1-2 h in the laminar tissue culture. Cells (1 × 10^6^/10 cm dish) were cultured in Dulbecco’s Modified Eagle Medium (DMEM; catalog no. 12-604F, Lonza) with 20% fetal bovine serum (FBS; catalog no. E5050-02, lot no. 70932, EURx), 0.1% fungizone, and 1% penicillin/streptomycin. Different lots of FBS from the same suppliers may vary and can affect AChR clustering. Therefore, testing different lots of FBS and selecting the one that efficiently facilitates the clustering of postsynaptic machinery are important. At all stages of culturing, the cells were evenly spread on the dish to prevent dedifferentiation. Cells were passaged every 3 days and before they reached 30% confluence (see Fig. 2 for recommended densities). After trypsinization, the cells were collected by centrifugation at 1700 rotations per minute for 3 min, resuspended in fresh culture medium, and split according to a 1:6 ratio onto fresh gelatin-coated dishes that contained warm culture medium. Cells that are passaged this way take ∼2 days to reach 30% confluence.

### Freezing C2C12 cells

Each 10-cm dish that contained C2C12 myoblasts was split between two freezing vials. After trypsinization, the cells were resuspended in 0.5 ml/vial of freezing medium I (DMEM with glutamine and 20% FBS at room temperature) and aliquoted into two separate vials. Pre-chilled freezing medium II (0.5 ml; DMEM with glutamine, 20% FBS, and 14% dimethylsulfoxide [DMSO; catalog no. A994.1, Roth]) was added. The vials were placed in a pre-chilled freezing container and moved immediately to −80°C. After 1-2 days, frozen cell stocks were placed in a liquid nitrogen container that was suitable for long-term storage.

### Thawing frozen cells

To prevent cell damage that could be caused by DMSO, the cells were thawed quickly in a 37°C water bath and immediately placed on a gelatin-coated dish that contained 10 ml of pre-warmed culture medium. The medium that contained remnants of the freezing medium was replaced the next day.

### Differentiation into myotubes

For myoblast fusion, we routinely use Permanox slides (catalog no. 160005, Thermo Fisher Scientific) and reusable Flexiperm eight-well grids (0.9 cm^2^ surface area/well), which is a suitable format for immunocytochemical microscopy and small-scale biochemical analysis. Flexiperm grids (catalog no. 6032039, Sarstedt) were sterilized by dipping in 100% ethanol, dried, and attached to Permanox slides. To induce AChR cluster formation, the Permanox surface was covered with 2 μg/ml laminin (catalog no. L2020-1MG, Sigma, and catalog no. LNKT-0201, Biolamina) in DMEM (200 μl/well). Frozen laminins were slowly thawed on ice to avoid aggregation that can result in uneven distribution on the slide. Importantly, the laminin solution needs to cover the well area evenly because uneven coating can affect the formation of AChR clusters and lead to the detachment of myotubes during differentiation. The Permanox slides were incubated with laminin overnight at 37°C in the tissue culture incubator. The next day, 1 × 10^5^ myoblasts were seeded per Flexiperm well (cell density of 1.4-1.6 × 10^5^/ml). Myoblast differentiation into myotubes was induced 24 h after cell seeding on laminin-coated Permanox slides because a longer incubation time can result in the degradation of laminin, which would affect AChR cluster formation. Myotube differentiation was induced by gentle replacement of the culture medium with fusion medium (DMEM with glutamine, 2% horse serum [HS], 0.1% fungizone, and 1% penicillin/streptomycin). Importantly, a substantial volume of the medium (∼700 μl/Flexiperm well) had to be added because replacing media during differentiation is not recommended. The cells were allowed to fuse for 4 days, with no cell agitation during that time because moving the slide or removing it from the incubator for microscopic inspection or medium replacement can significantly affect the attachment of large myotubes and interfere with their ability to form complex AChR clusters.

### Fixation and fluorescent staining

For fixation, the Permanox slides that contained differentiated myotubes were removed from the incubator and placed on a heat block at 37°C. The fusion medium was gently replaced with 4% paraformaldehyde (PFA) in phosphate-buffered saline (PBS). After the addition of PFA, the slides were removed from the heat block, followed by fixation for 7 min at room temperature. Paraformaldehyde was then removed, and each well was washed three times with PBS. For immunocytochemistry, the cells were blocked with blocking buffer (2% bovine serum albumin [BSA; catalog no. SC-2323, ChemCruz], 0.2% normal goat serum [NGS], and 0.05% Triton-X 100 [catalog no. T8787-100ML, Sigma]) for 30 min at room temperature. To visualize AChR clusters, F-actin cells were incubated for 30 min with AlexaFluor 488-α-bungarotoxin (1 ng/μl; catalog no. B13422, Invitrogen) and 100 nM Actistain conjugated to AlexaFluor 555 (in blocking buffer; catalog no. PHDH1, Cytoskeleton), followed by three washes with PBS. The Flexiperm grid was then gently removed, and 120 μl of Fluoromount was added to the Permanox slide, which was then sealed with a coverslip. For vinculin staining, primary antibody against vinculin (catalog no. V4506, Sigma) was used. For Ll5β staining, a previously described antibody was used^41^.

### Microscopic analysis

Microscopic analysis was performed at the Confocal Microscopy Facility, Nencki Institute, using a Zeiss Spinning Disc confocal microscope or Leica TCS SP8 scanning confocal microscope that was equipped with a diode or white light laser, respectively. Images were collected using ZEN software (ZEISS International) and analyzed using ImageJ/Fiji software.

### Silver staining

For silver staining analysis, Flexiperm wells were incubated with laminins overnight. The medium was then aspirated and 30 μl of 2× sample buffer (62.5 mM Tris-HCl [pH 6.8], 2.5% sodium dodecyl sulfate [SDS], 0.002% bromophenol blue, 0.7135 M [5%] β-mercaptoethanol, and 10% glycerol) that was added to each well. The solution was pipetted up and down 2-4 times, and the samples were then boiled at 95°C for 5 min. Proteins were resolved by SDS-polyacrylamide electrophoresis (PAGE). The gels were silver-stained according to the standard protocol.

### Western blot

The cells were lysed in lysis buffer (50 mM Tris-HCl, 150 mM NaCl, 1% Nonidet-P40, 0.5% SDS, 10% glycerol, 1 mM dithiothreitol [DTT], 1 mM NaF, and ethylenediaminetetraacetic acid-free mini protease inhibitor cocktail [catalog no. 11873580001, Roche], pH 8.0) and scraped off the dish. The lysates were briefly incubated on ice, passed three times through a 25-gauge needle, incubated on ice for 15 min, and centrifuged at 20,000 × *g* for 30 min at 4°C. For Western blot, the supernatant samples were mixed with sample buffer and boiled for 5 min. The proteins were then resolved by SDS-PAGE and transferred to nitrocellulose membranes (catalog no. 66485, Pall Corporation) using Trans Blot Turbo (catalog no. 1704270, Bio-Rad). Membranes were blocked with 5% milk in TBST (20 mM Tris-HCl, 150 mM NaCl [pH 7.6], and 0.1% Tween20) and probed with anti-AChR-α1 antibody (catalog no. 10613-1-AP, Proteintech) or anti-tubulin antibody (catalog no. ab18251, Abcam) in 5% milk in TBST. After washing with TBST buffer, the membranes were incubated with appropriate secondary antibodies conjugated to horseradish peroxidase. For protein detection, we used Clarity chemiluminescent substrate (catalog no. 1705060, Bio-Rad).

### Culturing human primary myoblasts

Human primary myoblasts were obtained from EuroBioBank. Cells were plated in F-10 Ham nutrient mixture (catalog no. N6908-500ML, Sigma) that contained L-glutamine, sodium bicarbonate, 20% FBS, 0.1% fungizone, and 1% penicillin/streptomycin. The medium was supplemented with 0.5 mg/ml fetuin (catalog no. F3385-100MG, Sigma), 0.39 μg/ml dexametazon (catalog no. D4902-25MG, Sigma), and 20 ng/ml basic fibroblast growth factor (catalog no. G507A, Promega). Human primary myoblasts were passaged the same way as C2C12 cells, with the exception of using 0.7% gelatin for coating the culture plates. Fusion was initiated 2 days after seeding 1 × 10^5^ cells/Flexiperm well by replacing the culture medium with 2% HS in DMEM (the same as for C2C12). The cells were allowed uninterrupted fusion for 12 days, after which they were fixed for microscopy.

## Data availability

The datasets generated during and/or analysed during the current study are available from the corresponding author on a reasonable request.

## Acknowledgements

We thank Dr. Marta Gawor for providing an image of AChR clusters in myotubes that were grown on agrin-coated substratum. This work was supported by Polish National Science Center Opus grant no. UMO-2018/29/B/NZ3/02675 (awarded to T.J.P.) and Preludium grant no. UMO-2018/29/N/NZ4/02724 (awarded to M.P). M.P was also supported by the European Union Horizon 2020 research and innovation program under Marie Sklodowska-Curie grant no. 665735 (Bio4Med) and the Polish Ministry of Science and Higher Education within 2016-2020 funds for the implementation of international projects (agreement no. 3548/H2020/COFUND/2016/2). The project was performed using the CePT infrastructure, financed by the European Union, European Regional Development Fund, within the Operational Program “Innovative Economy” for 2007-2013.

## Author contributions

M.P. designed and carried out the experiments and analyzed the data included in this publication. P.D. carried out the preliminary experiments. H.L. provided the primary human muscle cell line, analyzed the data and reviewed the manuscript. B.S.P supervised the scientific staff and edited the manuscript and figures. T.J.P. designed the experiments, analyzed the data, and provided scientific supervision. M.P. and T.J.P. wrote and reviewed the manuscript.

## Competing interests

We declare no competing interests.

## Figure Legends

**Supplementary Figure S1. Equal amounts of laminins were detected in the culturing wells after the coating procedure.** (**A**) Control for coating efficiency. After coating with each laminin, the wells were covered with sample buffer, and the collected material was resolved by SDS-PAGE and analyzed by silver staining. The results from three independent experiments are shown. (**B**) Predicted molecular weights for each laminin chain.

