## Supplementary figures and images for "An improved method for culturing myotubes on laminins for the robust clustering of postsynaptic machinery"

### Supplementary Figure 1

#### Supplementary figure 1.

**A**

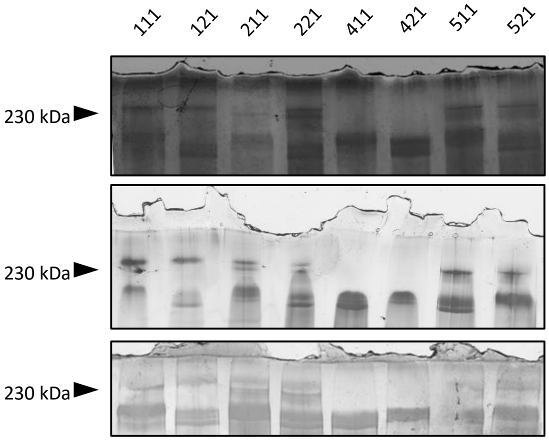


**B**

| Laminin chain | MW (kDa) |
|---|---|
| α1 | 337 |
| α2 | 343 |
| α4 | 203 |
| α5 | 400 |
| β1 | 198 |
| β2 | 196 |
| γ1 | 178 |
